# The clouded mother-of-pearl *Protogoniomorpha anacardii* Linnaeus, 1758 (Lepidoptera: Nymphalidae) – an updated record of butterfly for Saudi Arabia

**DOI:** 10.1101/2023.12.22.572884

**Authors:** Ateah Alfakih, Eisa Ali Al Faify

## Abstract

The record of the clouded mother-of-pearl *Protogoniomorpha anacardii* Linnaeus, 1758 (Lepidoptera: Nymphalidae) was updated in Saudi Arabia. At this stage, we confirmed the presence of this species in the southernmost part of Saudi Arabia based on photographs taken between 2020–2023 from Fayfa and the adjacent mountains in Jazan. The photographs match the descriptions and remarks made by Dr Torben Larsen in multiple publications, which suggest that this taxon represents the widespread subspecies *P. a. nebulosa* Trimen, 1881. Here, we discuss the Saudi Arabian populations of *P. a. nebulosa* and its conservation in depth and encourage more research on butterflies in Saudi Arabia.

## Introduction

The butterflies of Arabia have attracted many scientists and voyagers over the centuries. This can be seen in scientific records dating back to the beginning of the 19th century, when the 1820–1825 collections of Hemprich and Ehrenberg were presented in Klug’s works in 1829– 1845 (1-4). It is believed that the collection of the Arabian Rhopalocera had started before Hemprich and Ehrenberg’s travels since the Danish expedition to Arabia in 1761–1767, but the animal materials must have been damaged before they could be examined, unlike the plant materials (detailed in (2, 3)).

The most significant contribution to the butterflies of Saudi Arabia is credited to Dr Torben Larsen, who devoted a considerable part of his life to studying the butterflies of Arabia and discovered much of what we know today about them. Larsen’s journey in Arabia started in the 1970s and generated some publications on the Arabian butterflies, including from Oman in 1977 and 1980 (5-7), Saudi Arabia in 1979 (8), Yemen in 1982 (2) and then again from Saudi Arabia and its neighbours in 1984 (9). However, Larsen’s most comprehensive work was published as a monograph in 1983 (3), in which he thoroughly reviewed past studies on the butterflies of Arabia and raised the number of recorded butterflies in Arabia to more than 150 species. At this point (i.e. 1983), 24 butterflies of the Nymphalidae family were recorded in Saudi Arabia. This family is better characterised by the open cells in the hindwing (10) and the highly reduced forelegs compared to members of other families (9-11), which superficially look as if they possess only four legs. However, the absent native Arabian nymphalids from Saudi Arabia belong to the three genera of *Neptis* Fabricius, 1807, *Phalanta* Horsfield, 1829 and *Protogoniomorpha* Wallengren, 1857, recorded only in Yemen (9, 12). Later, Pittaway (13) studied the butterfly diversity in the western part of Saudi Arabia and published a book with Walker (14). Still, these works failed to record the nymphalids belonging to the above three genera. To the best of our knowledge, the records of additional genera of butterflies in Saudi Arabia have not much advanced beyond the last recording of an alien species from the eastern part of the country (15). However, during the preparation of this article, Tshikolovets was able to document new genera and species of butterflies for Saudi Arabia that includes the clouded mother-of-pearl *Protogoniomorpha anacardii* Linnaeus, 1758 (16). Here, we provide an updated record of this butterfly from Saudi Arabia with an expanded discussion on its ecology and conservation.

## Methods

Due to the rarity of the clouded mother-of-pearl *P. anacardii* butterfly, we did not use invasive methods, such as netting or bait trapping, to collect specimens for description purposes. Thus, for the description in this article, we relied on images. As this species is the only one in Arabia from the genus *Protogoniomorpha*, reliance on photographs will not cause misidentification. In the summer of 2020, E.A. Al Faify captured the first-ever photograph of the clouded mother-of-pearl from Fayfa, Saudi Arabia. However, due to the coronavirus disease 2019 pandemic, we delayed the work on this record until the restrictions were lifted, allowing us to collect some samples for description purposes. However, it is still extremely difficult to find this butterfly in great abundance. In January 2021, E.A. Al Faify took another photograph of this butterfly from the same locality. The butterfly’s presence was confirmed in February and May 2023 with a pattern of abundance in the same locality and adjacent mountains, raising the observations to more than 10 during the past three years (Figure 1; Table 1). However, we preferred not to use traps until we obtained more information about the distribution pattern in the Jazan mountains.

**Table 1:** The localities and dates of *P. anacardii*’s current record from Saudi Arabia.

| Locality | Coordinates | Altitude | Year |
| --- | --- | --- | --- |
| Fayfa | 17°14'54.3"N 43°05'48.1"E | 1300m | 2020 and 2023 |
| Fayfa | 17°15'16.9"N 43°04'31.7"E | 980m | 2020 |
| Fayfa | 17°15'36.6"N 43°05'52.2"E | 1280m | 2021 |
| Fayfa | 17°15'37.1"N 43°05'58.1"E | 1360m | 2021 |
| Bani Malik | 17°18'41.4"N 43°09'21.0"E | 1050m | 2020 |

**Figure 1:**
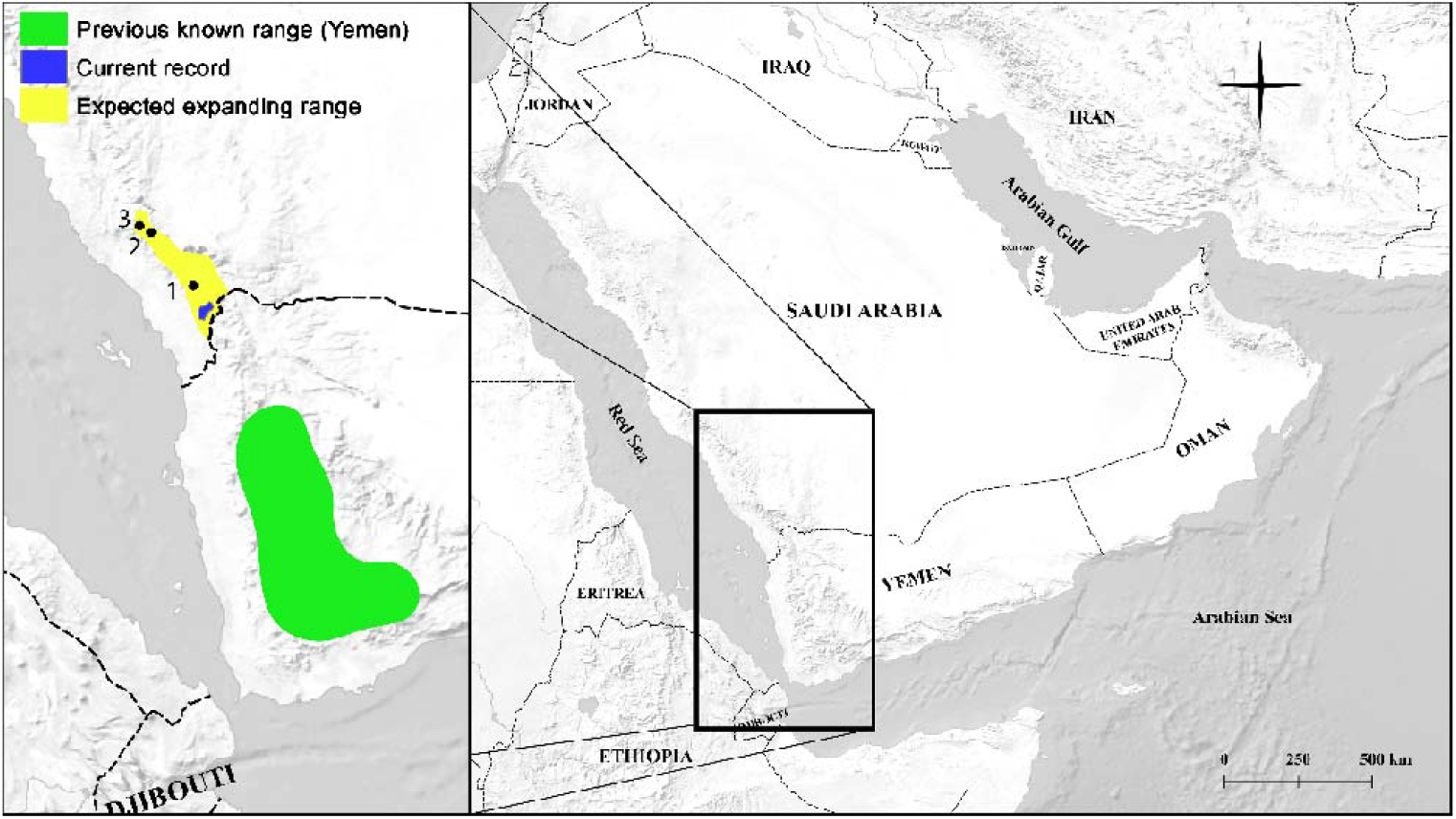
Distribution of *P. anacardii* in Arabia. The green overlay depicts the approximate previous records in Yemen (reported by (2, 3, 17)). The range of the current records in Saudi Arabia is highlighted in blue, which approximately coincides with Tshikolovets’s record (16), while the predicted expanding range is in yellow (1: Harub mountains, 2: Raidah Escarpment and 3: Rijal Almaa).

## Results and Discussion

*Protogoniomorpha* is an Afrotropical genus that contains a few species (11, 18) distributed in Africa, with only one species, *P. anacardii*, which extends to Arabia in Yemen: (2, 3, 9, 12, 17) and Saudi Arabia (16). This species is further classified into four subspecies (18-20): *P. a. anacardii* Linnaeus, 1758 in West Africa, *P. a. ansorgei* Rothschild, 1904 in Angola and the Democratic Republic of Congo, *P. a. duprei* Vinson, 1863 in Madagascar and the widely distributed *P. a. nebulosa* Trimen, 1881, which extends from South Africa to East Africa, alongside the countries adjacent to the rift valley, and is restricted in Yemen from Arabia (2, 3, 9, 12, 17, 18, 21). In Africa, this species possesses two different seasonal forms: wet and dry (11, 22). However, in Arabia, it has displayed only a pale form to date (see below). In this article, it is premature to determine the subspecies level because we did not collect specimens and had no access to preserved specimens. Although Larsen adhered to Gabriel’s (17) identification of the subspecies *P. a. nebulosa*, he was conservative in determining the subspecies level in Arabia (3) and even in Africa (21). Recently, Tshikolovets regarded it as *P. a. nebulosa* (16), and thus, we adhere to the identification of the subspecies *P. a. nebulosa* to prevent unnecessary inconsistencies. Still, further research is needed to determine whether this butterfly represents an independent subspecies.

The photographed butterfly (Figure 2A–D) cannot be mistaken for other species from Arabia or Africa. It can be readily identified as *P. anacardii* due to the extensive black markings in the wing apex and margins (10, 21), which appear to be a stable trait. The characteristics that appeared in the photographs (Figure 2A–D), with emphasis on the pale form, match the descriptions and figures in: Larsen’s works (2, 3, 9, 18), figures in Tshikolovets’s work (16) and description in Gabriel’s work (17). A literature search revealed that this butterfly was recorded for the first time in Arabia from Yemen (17) after collections were made by Scott and Britton during the British Natural History Museum’s southwest Arabian expedition from 1937 to 1938. At the time of the first record, Gabriel (17) identified several butterfly species from Saudi Arabia collected by John Philby in 1936 from Fayfa and Asir mountains, but Philby failed to record *P. anacardii*.

**Figure 2.**
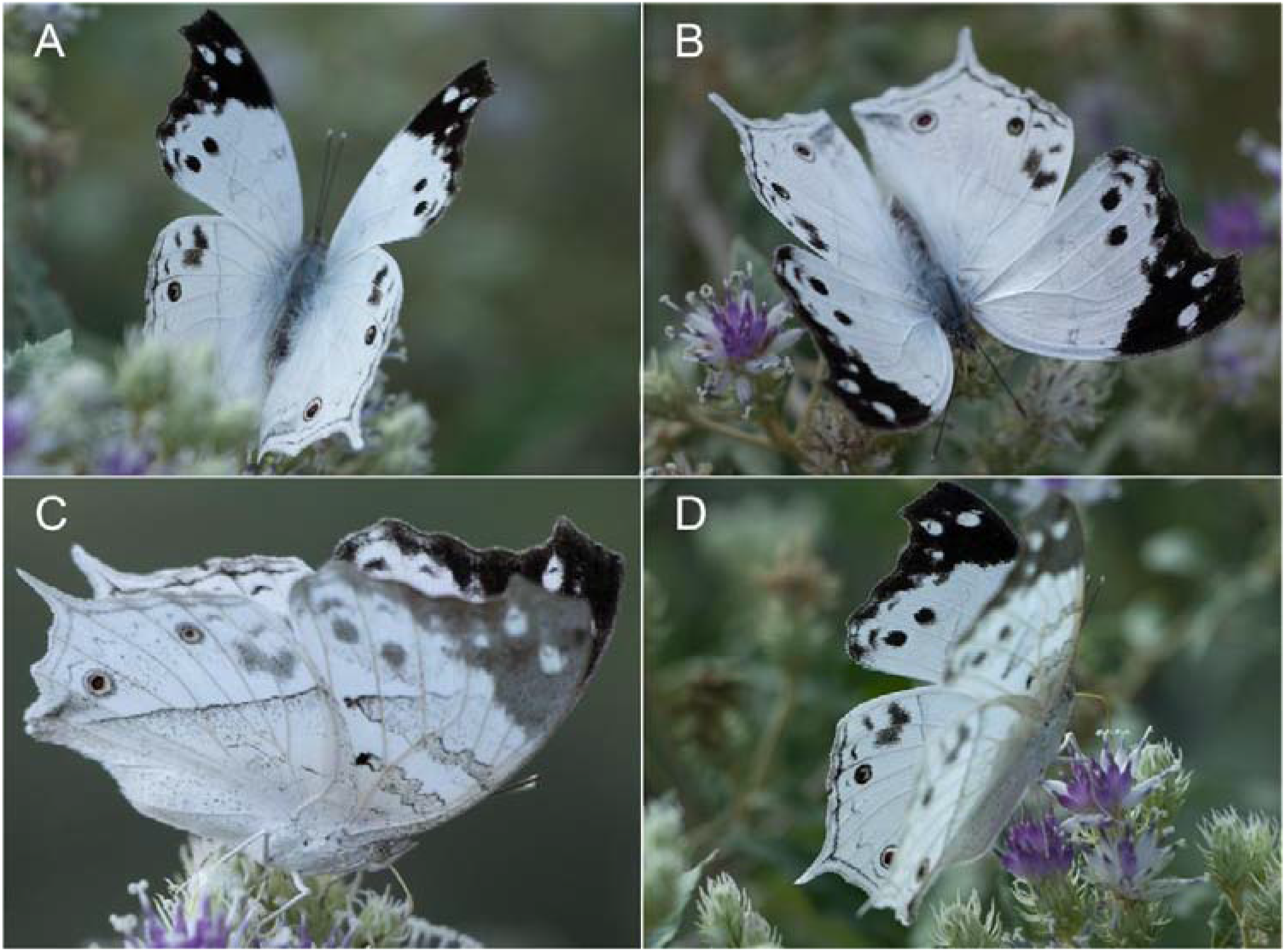
**A–D:** An individual of *P. anacardii* on *Baccharoides schimperi* (DC.) Isawumi, El-Ghazaly & B. Nord. (Asteraceae), displaying the upper and undersides from Fayfa, Saudi Arabia (Photos: E.A. Al Faify).

The clouded mother-of-pearl was believed to have been of a recent entry into Arabia (17). However, from the literature presented above, Larsen must not have overlooked this large species during his extensive Rhopalocera collection journeys in Saudi Arabia, but he expected the spread of more species from Yemen northward to Saudi Arabia. Larsen (3) clearly stated that:

> Some of the known Arabian species will be found to have rather more extensive distribution areas than is known at present. Many Afrotropical species currently known only from Yemen and PDRY [People’s Democratic Republic of Yemen] will eventually be found in the Asir and the Hadhramaut, some possibly even in the Hejaz or northern Oman. 3(p.342).

The over 10 sightings of the butterfly within the past three years in this record and in 2021 by Tshikolovets (16) suggest that it might have only recently entered Saudi Arabia. However, the ecological parameters in the eastern Jazan mountains match those in the Yemeni habitats (23), where the butterfly thrives (see (2, 17) for details about the habitats’ descriptions). Sometimes, this species is observed in gregarious individuals in Africa (10, 21), which may indicate migration (21). This butterfly probably migrates to Saudi Arabia from time to time. For example, the sooty orange tip, *Zegris eupheme larseni* Pittaway, 1985, was collected in large numbers in 1983 in Haql, north of NEOM, but it was completely absent in the subsequent year (13). Therefore, we need records of early life stages, including eggs, larvae and pupae, to confirm whether viable breeding populations have been established and to predict the dispersal in Saudi Arabia.

In Africa, the clouded mother-of-pearl is reported to occupy forest margins, coastal areas and savannah woods (10) from sea level up to 2000 m, especially in areas dominated by the trees of *Brachystegia* Benth. (11). Its larvae generally feed on several Acanthaceae plants (11, 18, 21, 22). In Yemen, it has been reported from only the southernmost part of the country in riverine forests from 900 m to 1500 m sea levels (2, 3). These areas are characterised by high precipitation, dense vegetation and agricultural terraces.

The localities of the records in this study (Table 1) represent plant diversity hotspots in Saudi Arabia (24), which are likely suitable habitats for the butterfly to establish viable breeding populations and extend its range northward. Fayfa and the close mountains benefit from the relatively high precipitation, which allows sustained vegetation cover. Thirteen genera of the Acanthaceae family, as shown in Table 2, are present in Fayfa and its neighbouring areas, comprising 25 species (23, 25-27).

**Table 2:** Genera and species count of the Acanthaceae family from Fayfa and its neighbouring areas. Incorporated from (23, 25-27).

| Genus | Species |
| --- | --- |
| *Anisotes* | *trisulcus* (Forssk.) Nees |
| *Asystasia* | *gangetica* (L.) T.Anderson |
| *Barleria* | *acanthoides* Vahl<br>*bispinosa* (Forssk.) Vahl |
|  | *hochstetteri* Nees<br>*prionitis* L.<br>*proxima* Lindau<br>*trispinosa* (Forssk.) Vahl |
| *Blepharis* | *edulis* (Forssk.) Pers. [Sometimes misidentified for *B. ciliaris* (L.)<br>B.L.Burt] |
|  | *maderaspatensis* (L.) B.Heyne ex Roth |
| *Crossandra* | *johanninae* Fiori |
| *Dicliptera* | *paniculata* (Forssk.) I.Darbysh. |
| *Ecbolium* | *gymnostachyum* (Nees) Milne-Redh.<br>*viride* (Forssk.) Alston |
| *Hypoestes* | *forskaolii* (Vahl) R.Br. |
| *Justicia* | *caerulea* Forssk.<br>*flava* (Forssk.) Vahl<br>*heterocarpa* T.Anderson<br>*odora* (Forssk.) Lam. |
| *Lepidagathis* | *scariosa* Nees |
| *Meiosperma* | *debile* (Forssk.) I.Darbysh. & E.A.Tripp |
| *Phaulopsis* | *imbricata* (Forssk.) Sweet |
| *Ruellia* | *forsskaolii* Thulin<br>*patula* Jacq.<br>*prostrata* Poir. |

Eight of these species are the endangered *Asystasia gangetica, Barleria prionitis, B. proxima, Blepharis maderaspatensis, Lepidagathis scariosa, Justicia caerulea, Meiosperma debile* and *Phaulopsis imbricata* (23, 25-27), with *J. caerulea* and *P. imbricata* seen only in Fayfa and its very close vicinity in Saudi Arabia (26, 27). The most reported host plants that *P. anacardii*’s larvae feed on in Africa are *Asystasia spp*. including *A. gangetica* (11, 18, 21, 22). In Saudi Arabia, this plant can be found at lower elevations (about 250 m) and drier lands, seemingly in only one proximate location to the locations of the current records (Table: 1)—Al Aridhah (25-27). Indeed, this butterfly was observed in dry wadis with a few trees in Yemen (2, 3). Therefore, recording this species in drier and lower lands remains probable. Although the larvae seem to be generalists feeding on multiple species, it is challenging to determine whether the presence of the clouded mother-of-pearl will exert pressure on any of the aforementioned endangered plants. Therefore, we recommend a reassessment of these endangered species and whether *P. anacardii* exploits them.

As observed in their African counterparts (10, 22), adult butterflies in Fayfa were observed flying in summer and winter. Moreover, they were seen to visit flowers of the Asteraceae family (Figure 2A–D) in winter and perch on and hide beneath cultivated banana trees during the day and night, which confirms the previous observations in Yemen that may indicate the preference of the butterfly for banana terraces (17). It is acknowledged that this species is Pan-African, meaning it has a wide ecological tolerance (12). Even though dry lands separate the southwestern mountains in Saudi Arabia (24), a few discrete escarpments lying within those dry areas are more humid and densely vegetated. Therefore, based on the habitat parameters of the records from Saudi Arabia in the present study and Yemen (2, 17), populations of *P. anacardii* may eventually spread northward to the Harub mountains in the Jazan region, Rijal Almaa and Raidah Escarpment in the Asir region (Figure 1). Above all, it has been nearly four decades since the last discovery of native butterflies in Saudi Arabia, and the work on butterflies needs to be updated. We strongly uphold Larsen’s (3) and Pittaway’s (13) views, suggesting the existence of newer species in Saudi Arabia’s southern and northern mountains. We encourage researchers to explore Saudi Arabia’s northernmost areas, including NEOM, and compare the Rhopalocera species with the Levant and Egyptian faunas, which have received less attention than the southern mountains.

## Acknowledgements

We thank Ali Alzahrani, Nafee Alothayqi and Saad Albaqami for their valuable comments on the manuscript.

## References

1. Baker DB. C. G. Ehrenberg and W. F. Hemprich’s travels, 1820–1825, and the Insecta of the Symbolae Physicae. Deutsche Entomologische Zeitschrift. 1997;44(2):165–202.

2. Larsen TB. The butterflies of the Yemen Arab Republic (with a review of the species in the Charaxes viola-group from Arabia and East Africa). Biologiske Skrifter, Kongelige Danske Videnskabernes Selskab. 1982;23(3):1–87.

3. Larsen TB. Insects of Saudi Arabia. Lepidoptera; Rhopalocera (a monograph of the butterflies of the Arabian Peninsula). Fauna of Saudi Arabia. 1983;5:333–478.

4. Olivier A, Nekrutenko YP. The butterflies described by Johann Christoph Friedrich Klug (1775–1856) in his Symbolae Physicae, Insecta (Lepidoptera, Pieridae, Lycaenidae, Nymphalidae): An annotated review, with a catalogue of the existing types. Deutsche Entomologische Zeitschrift. 2000;47(1):95–104.

5. Larsen K, Larsen TB. Butterflies of Oman. Edinburgh: John Bartholomew & Son. Ltd.; 1980.

6. Larsen TB. The butterflies of eastern Oman and their zoogeographic composition. In: The scientific results of the Oman flora and fauna survey 1975. Journal Of Oman Studies, Special Report. 1977;1:179–208.

7. Larsen TB. The butterflies of Dhofar and their zoogeographic composition. In: The scientific results of the Oman flora and fauna survey 1977 (Dhofar). Journal Of Oman Studies, Special Report. 1980;2:153–186.

8. Larsen TB. Insects of Saudi Arabia. Lepidoptera: fam. Papilionidae, Pieridae, Danaidae, Nymphalidae, Lycaenidae. Fauna of Saudi Arabia. 1979;1:342–344.

9. Larsen TB. Butterflies of Saudi Arabia and Its neighbours. London: Stacey International; 1984.

10. Williams JG. A Field Guide to the Butterflies of Africa. London: Collins; 1969.

11. Kielland J. Butterflies of Tanzania. Melbourne: Hill House; 1990.

12. Larsen TB. The zoogeographical composition and distribution of the Arabian butterflies (Lepidoptera; Rhopalocera). Journal of Biogeography. 1984;11(2):119–158.

13. Pittaway AR. Lepidoptera: Rhopalocera of western Saudi Arabia. Fauna of Saudi Arabia. 1985;7:172–197.

14. Walker DH, Pittaway AR. Insects of Eastern Arabia. London: Macmillan Publishers Ltd.; 1987.

15. Pittaway T, Larsen TB, Legrain A, Majer J, Weidenhoffer Z. The establishment of an American butterfly in the Arabian Gulf: Brephidium exilis (Boisduval, 1852) (Lycaenidae). Nota Lepidopterologica. 2006;29(1/2):5–16.

16. Tshikolovets V. Notes on butterflies of Saudi Arabia with a description of a new subspecies (Rhopalocera, Lycaenidae, Nymphalidae et Hesperiidae). Atalanta. 2022;53(1/2): 205–208.

17. Gabriel AG. Lepidoptera Rhopalocera. British Museum (Natural History) Expedition to South-West Arabia, 1937-8. 1954;1(25): 351–391.

18. Larsen TB. Butterflies of West Africa. Stenstrup: Apollo Books; 2005.

19. Rothschild W, Jordan K. Lepidoptera collected by Oscar Neumann in North-East Africa. Novitates Zoologicae: A Journal of Zoology in Connection with the Tring Museum. 1903;10:491–542.

20. Rothschild W. New forms of butterflies. Novitates zoologicae: A Journal of Zoology in Connection with the Tring Museum. 1904;11:452–455.

21. Larsen TB. The butterflies of Kenya: and their natural history. New York: Oxford University Press; 1991.

22. Migdoll I. Field guide to the butterflies of Southern Africa. Cape Town: Struik Publishers; 1994.

23. Al-Turki TA. A prelude to the study of the flora of Jabal Fayfa in Saudi Arabia. Kuwait Journal of Science & Engineering. 2004;31(2):77–145.

24. Miller AG, Nyberg JA. Patterns of endemism in Arabia. Flora et Vegetation Mundi. 1991;9:263–279.

25. Collenette IS. An illustrated guide to the flowers of Saudi Arabia. London: Scorpion Publishing Ltd.; 1985.

26. Collenette IS. Wildflowers of Saudi Arabia. Riyadh: National Commission for Wildlife Conservation and Development (NCWCD); 1999.

27. Alfarhan AH, Al-Turki TA, Basahy AY. Flora of Jizan region, final report of project AR-17-7. Saudi Arabia: King Abdulaziz City for Science and Technology; 2005.

